# A SARS-CoV-2 Vaccination Strategy Focused on Population-Scale Immunity

**DOI:** 10.1101/2020.03.31.018978

**Authors:** Mark Yarmarkovich, John M. Warrington, Alvin Farrel, John M. Maris

**Affiliations:** Division of Oncology and Center for Childhood Cancer Research; Children’s Hospital of Philadelphia; Philadelphia, PA, 19104; USA; Department of Biomedical and Health Informatics, Children’s Hospital of Philadelphia; Philadelphia, PA, 19104; Perelman School of Medicine at the University of Pennsylvania; Philadelphia, PA, 19104

**Author notes:** **Corresponding author** John M. Maris, MD, The Children’s Hospital of Philadelphia, Colket Translational Research Building, Rm. 3030, 3501 Civic Center Blvd., Philadelphia, PA, 19104.

**Keywords:** COVID19, SARS-CoV-2, coronavirus, vaccine, DNA vaccine, RNA vaccine

## Abstract

Here we propose a vaccination strategy for SARS-CoV-2 based on identification of both highly conserved regions of the virus and newly acquired adaptations that are presented by MHC class I and II across the vast majority of the population, are highly dissimilar from the human proteome, and are predicted B cell epitopes. We present 65 peptide sequences that we expect to result in a safe and effective vaccine which can be rapidly tested in DNA, mRNA, or synthetic peptide constructs. These include epitopes that are contained within evolutionarily divergent regions of the spike protein reported to increase infectivity through increased binding to the ACE2 receptor, and within a novel furin cleavage site thought to increase membrane fusion. This vaccination strategy specifically targets unique vulnerabilities of SARS-CoV-2 and should engage a robust adaptive immune response in the vast majority of the human population.

## Introduction

The current SARS-CoV-2 pandemic has precipitated an urgent need to rapidly develop and deploy a safe and effective vaccine. Optimally designed vaccines maximize immunogenicity towards regions of proteins that contribute most to protective immunity, while minimizing the antigenic load contributed by unnecessary protein domains that may result in autoimmunity, reactogenicity, or even enhanced infectivity. Here we propose a vaccine concept that focuses on: 1) stimulation of CD4 and CD8 T cells, 2) immunogenicity across the majority of human HLA alleles, 3) targeting both evolutionarily conserved regions, as well as newly divergent regions of the virus that increase infectivity, 4) targeting linear and conformational B cell epitopes, and 5) targeting viral regions with the highest degree of dissimilarity to the self-immunopeptidome, maximizing safety and immunogenicity. We present a list of viral antigen minigenes for use in a multivalent vaccine construct that can be delivered by scalable techniques such as DNA, nucleoside mRNA, or synthetic peptides.

SARS-CoV-2 is the third coronavirus in the past two decades to acquire infectivity in humans and result in regional epidemics, with SARS-CoV-2 causing a global pandemic. The spike glycoprotein of SARS-CoV-2 dictates species tropism and is thought to bind to ACE2 receptors with 10-20-fold higher affinity than SARS-CoV in humans (Walls et al.; Wrapp et al., 2020). In addition, cleavage at a novel furin insertion site is predicted to facilitated membrane fusion and confer increased virulence, as has been previously reported with other viruses (Chen et al., 1998). Based on initial reports, infection of ACE2-expressing pneumocytes lining the pulmonary alveoli likely impairs release of surfactants that maintain surface tension, hindering the ability to prevent accumulation of fluid that may lead to acute respiratory distress syndrome (Xu et al., 2020; Zhang et al., 2020). The immune response of convalescent COVID-19 patients consists of antibody-secreting cells releasing IgG and IgM antibodies, increased follicular helper T cells, and activated CD4 and CD8 T cells (Thevarajan et al., 2020), suggesting that a broad humoral and T cell driven immune response mediates the clearance of infection. The large size of the SARS-CoV-2 (∼29kb) suggests that selection of optimal epitopes and reduction of unnecessary antigenic load for vaccination will be essential for safety and efficacy.

Rapid deployment of antibody-based vaccination against SARS-CoV-2 raises a major concern in accelerating infectivity through Antibody-Dependent Enhancement (ADE), the facilitation of viral entry into host cells mediated by subneutralizing antibodies (those capable of binding viral particles, but not neutralizing them) (Dejnirattisai et al., 2016). ADE mechanisms have been described with other members of the *Coronaviridae* family (Wan et al., 2020; Wang et al., 2016), and it has already been suggested that some of the heterogeneity in COVID-19 cases may be due to ADE from prior infection from other viruses in the coronavirus family (Tetro, 2020). We suggest that although the T cell epitopes presented here are expected to be safe in vaccination, B cell epitopes should be further evaluated for their ability to induce neutralizing antibodies as compared to their potential to induce ADE. As it has been shown that T helper (T_H_) cell responses are essential in humoral immune memory response (Alspach et al., 2019; McHeyzer-Williams, Okitsu, Wang, & McHeyzer-Williams, 2012), we expect that the T cell epitopes presented here will activate a CD4 T cells and drive memory B cell formation when paired with matched B cell epitopes.

The potential of a peptide-based vaccine to induce a memory B and T cell response is complicated by the diversity of HLA alleles across the human population. The HLA locus is the most polymorphic region of the human genome, resulting in differential presentation of antigens to the immune system in each individual. Therefore, individual epitopes may be presented in a mutually exclusive manner across individuals, confounding the ability to immunize with broadly presented antigens. While T cell receptors (TCRs) recognize linearized peptides anchored in the MHC groove, B cell receptors (BCRs) can recognize both linear and conformational epitopes, and are therefore difficult to predict without prior knowledge of a protein structure. Here we describe an approach for prioritizing viral epitopes and present a list of peptides predicted to safely target the vulnerabilities of SARS-CoV-2, generating highly immunogenic epitopes on both MHC class I and II in the vast majority of the population, increasing the likelihood that prioritized epitopes will drive an adaptive memory response.

## Results

We used our recently published methods for scoring population-scale HLA presentation of individual putative cancer antigens along the length of a protein to analyze the population-scale HLA presentation of individual peptides derived from all 10 SARS-CoV-2 genes across 84 Class I HLA alleles (Yarmarkovich et al., 2020), representing 99.4% of the population represented in the Bone Marrow Registry (Gragert, Madbouly, Freeman, & Maiers, 2013). We identified 3524 SARS-CoV-2 epitopes that are predicted to bind at least one HLA class I allele, with peptide FVNEFYAYL capable of binding 30 unique HLA alleles representing 90.2% of the US population (**Figure 1A, top; Table 1; Table S1**). We tested various epitope sizes to maximize HLA presentation across the viral proteome, finding that 33 amino acid epitopes generated maximal population-scale HLA presentation, and suggest that these 33mers can be expressed in a multicistronic construct in dendritic cells to induce potent immune response across the vast majority of the population (An, Rodriguez, Harkins, Zhang, & Whitton, 2000; Lu et al., 2014). We identified areas predicted to be presented across the majority of the population, including a single 33mer ISNSWLMWLIINLVQMAPISAMVRMYIFFASFY containing epitopes capable of binding 82 of the 84 HLAs alleles studied here (**Table S1B**).

**Table 1.**
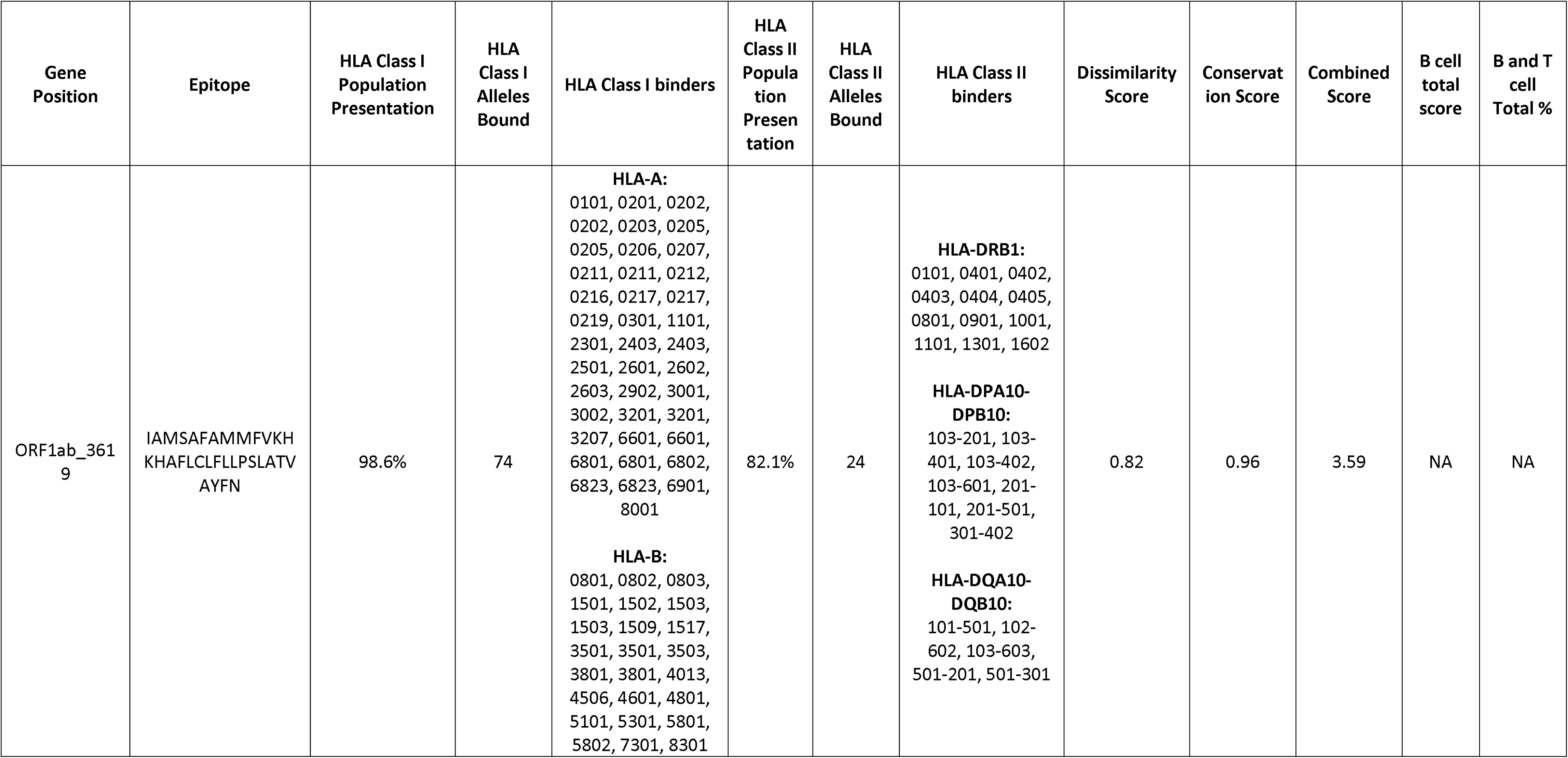
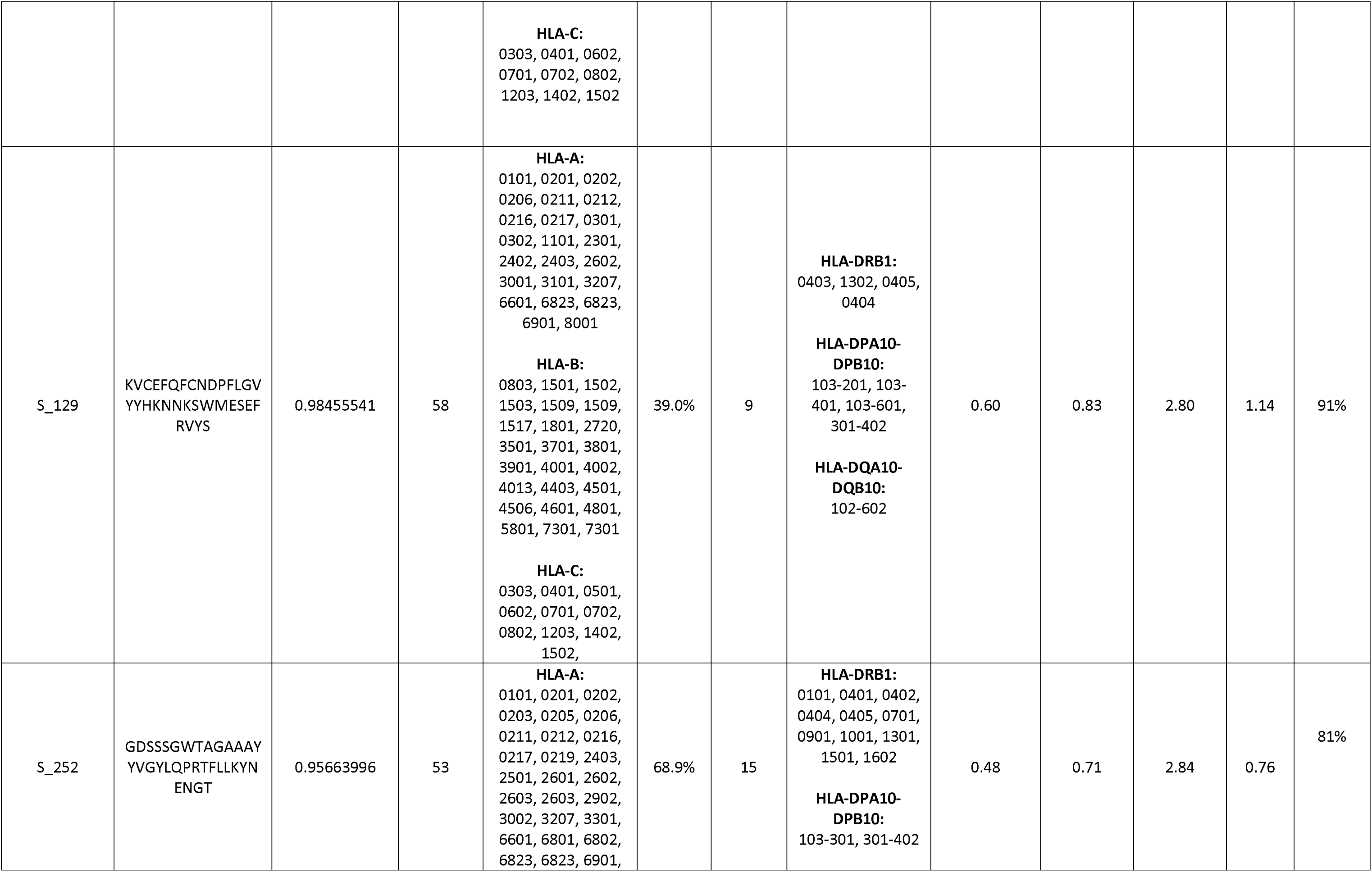
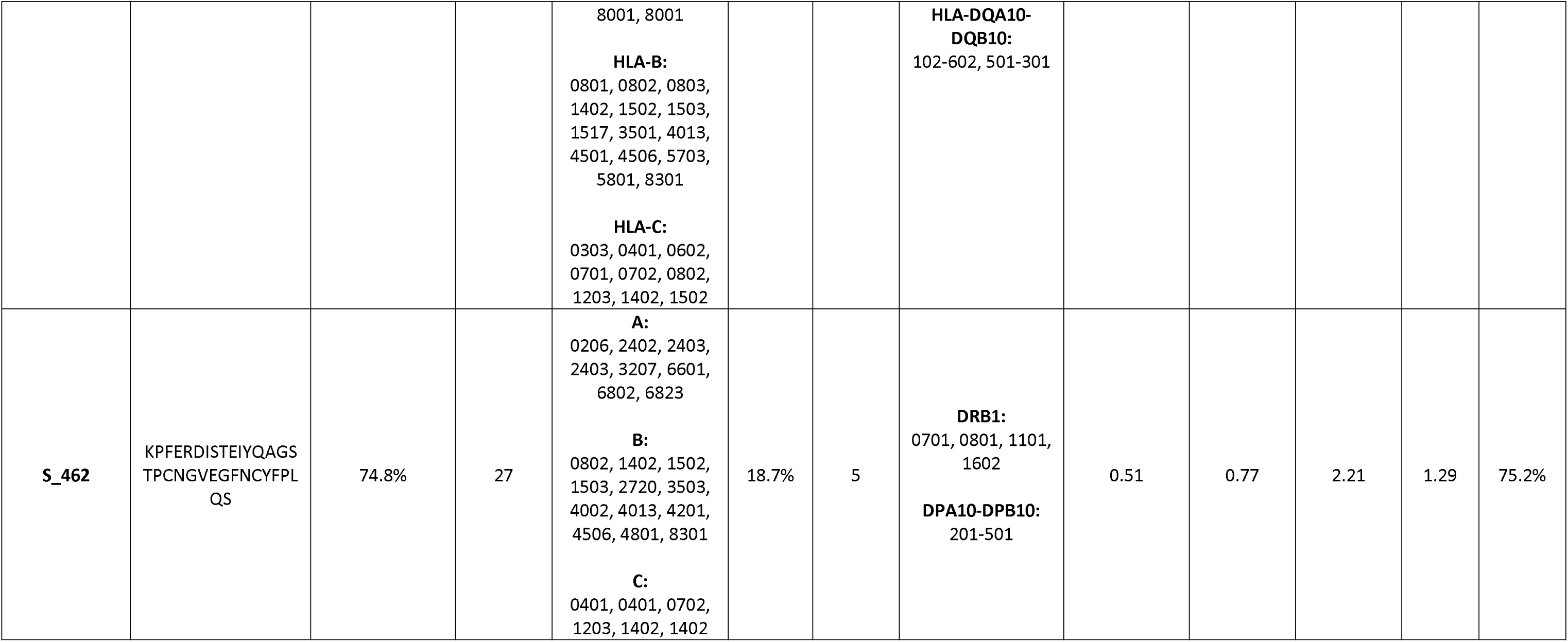
List of highest scoring viral epitopes suggested for vaccination based on MHC class I population-scale presentation, MHC class II population presentation, similarity score, and homology score across 9 mammal species and 1,024 human SARS-CoV-2 cases. Last column represents total score across all parameters, highlighting epitope S_462 in S protein containing novel receptor binding sites (Shang et al., 2020).

**Figure 1.**
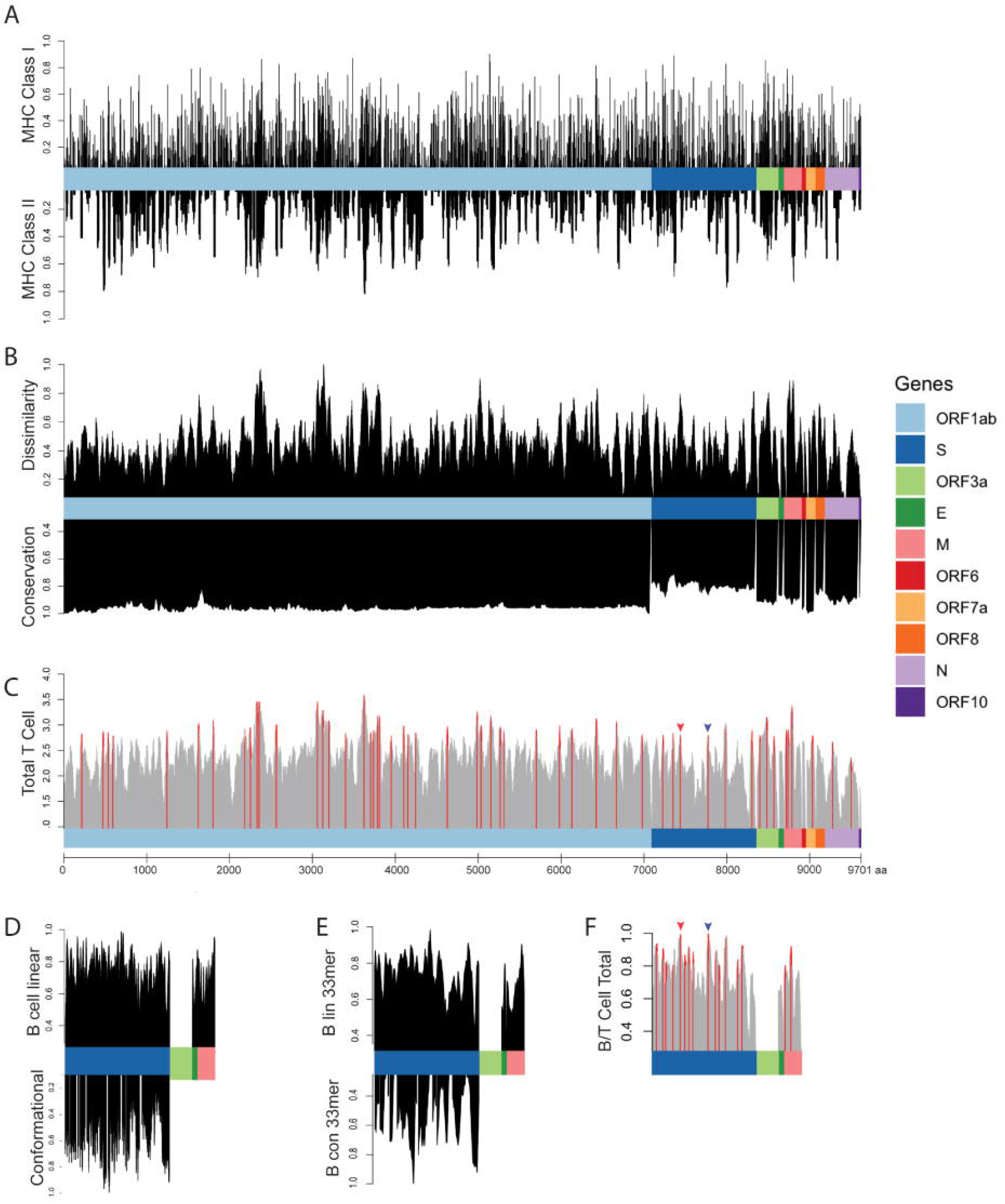
Epitope Scoring along SARS-CoV-2 Proteome. A) HLA presentation of 33mers across viral proteome. Representation of MHC Class I presentation (top) and MHC Class II presentation (bottom) reported as frequency of the population predicted to present each region of the viral proteome. B) Scoring of each epitopes along the length of the proteome as compared to the epitopes derived from the normal human proteome presented across 84 HLA alleles, reported as normalized scores in which the highest scoring epitopes are maximally dissimilar to self-peptides derived from normal proteins (top). Scoring for genomic conservation against 9 cross-species coronaviruses and 1,024 human sequences, with highest scoring regions conserved across human and other mammalian coronaviruses (bottom). C) Combined epitope score reported as sum of four above parameters (local maximum for epitopes with 90^th^ percentile total score in red). D) Scoring of B cell epitopes for each amino acid for linear epitopes in for Spike, Envelope, and Matrix proteins (top) and conformational epitopes in Spike protein (bottom). E) Combined scoring of 33mer epitopes as described in D. F) Combined B and T cell epitope scoring in Spike, Envelope, and Matrix proteins. Receptor binding domain epitope highlighted with red arrow and epitope containing furin cleavage site highlighted with blue arrow (**Figure 2**)

As it has been shown that presentation by both Class I and Class II MHC is necessary for robust memory B and T cell responses (Alspach et al., 2019; McHeyzer-Williams et al., 2012), we next analyzed presentation of these viral epitopes on 36 MHC Class II HLA alleles, representing 92.6% of the population (**Figure 1A, bottom; Table 1; Table S1**). Peptides derived from the 33mer IAMSAFAMMFVKHKHAFLCLFLLPSLATVAYFN were presented on 24 HLA class II alleles, representing 82.1% of US population, and peptides from the same epitope were predicted to be presented on 74 HLA class I alleles with a population frequency of 98.6%. As HLA frequencies vary based on the composition of each population, the frequency of individual HLA alleles can be adjusted based on specific populations using a SARs-CoV-2 immunogenicity map created here (**Table S1**).

Next, we sought to identify the most highly conserved regions of the SARS-CoV-2 virus, positing that non-conserved regions that are not involved in newly acquired increased infectivity may be prone to T cell evasion through mutation of MHC-presented epitopes. To do this, we compared the amino acid sequence of SARS-CoV-2 to fourteen *Coronaviridae* family sequences derived from bats, pigs, and camels, scoring each amino acid for conservation across the viral strains. We also scored the conservation across the 1,024 SARS-CoV-2 virus sequences available at the time of this analysis, equally weighing contributions from cross-species and interhuman variation (scores normalized to 0-1, with entirely conserved regions scoring 1). As expected, evolutionary divergence was greatest in the tropism-determining Spike protein and lowest in ORF1ab which contains 16 proteins involved in viral replication (**Figure 1B, bottom**).

We then compared predicted viral MHC-presented epitopes to self-peptides presented normally on 84 HLA alleles across the entire human proteome from UniProt, prioritizing antigens that are most dissimilar from self-peptides based on: 1) higher predicted safety based on less likelihood of inducing autoimmunity due to cross-reactivity with similar self-peptides presented on MHC; and 2) higher immunogenicity of dissimilar peptides based on an expected greater repertoire of antigen-specific T cells due to lower degree of negative thymic selection. To do this, we compared 3,524 viral epitopes against the normal human proteome on each of their MHC binding partners, testing a total of 12,383 peptide/MHC pairs against the entire human proteome (85,915,364 normal peptides across HLAs), assigning a similarity score for each peptide, with high scoring peptides representing the highest degree of dissimilarity as compared to the space of all possible MHC epitopes derived from the normal proteome (**Methods; Figure 1B, bottom; Table S1**).

To assign an overall score for T cell antigens, we normalized each of our four scoring parameters (represented in **Figure 1A and 1B**) between 0-1 and summed each metric to obtain a final epitope score, highlighting the local maxima of epitopes scoring in the 90^th^ percentile, highlighting 55 top scoring T cell epitopes across 9 SARS-CoV-2 genes as epitopes for vaccination (**Figure 1C red bars, Table S2**).

Finally, we sought to characterize B cell epitopes, assessing linear epitopes in Spike (S), Matrix (M), and Envelope (E) proteins which are exposed and expected to be accessible to antibodies, and characterized conformational epitopes in the Spike protein for which structural data are available using BepiPred 2.0 and DiscoTope 2.0 (Jespersen, Peters, Nielsen, & Marcatili, 2017; Kringelum, Lundegaard, Lund, & Nielsen, 2012). There was a strong concordance between linear epitope scores and conformational epitope scores (p<2e^-16^). We next performed an agnostic scoring of individual amino acid residues in S, M, and E proteins (**Figure 1D**), and then used these scores to generate scores for 33mer epitopes along the length of the protein (**Figure 1E**). The 33mer VGGNYNYLYRLFRKSNLKPFERDISTEIYQAGS derived from S protein at position 445 ranked the highest based on combined linear and conformational B cell epitope scoring. We combined T cell epitope scores calculated above with available B cell epitope scores derived from the S, M, and E genes, providing a list of antigens predicted to stimulate both humoral and cellular adaptive immunity (**Figure 1F**).

In addition to prioritizing evolutionarily conserved regions, we sought to specifically target acquired vulnerabilities in SARS-CoV-2 by focusing on novel features of this coronavirus that have been shown to contribute to its increased infectivity. The receptor binding domain of the SARS-CoV-2 Spike protein has been reported to have 10-fold higher binding affinity to ACE2 (Wrapp et al., 2020). We show that viral epitope GEVFNATRFASVYAWNRKRISNCVADYSVLYNS derived from the receptor binding domain (RBD) of the Spike protein (position 339-372) scores in the 90.9^th^ percentile of T epitopes and is the #3 of 1,546 epitopes scored in the S, E, and M genes for combined B and T cell epitopes, with presentation by MHC class I in 98.3% of the population (**Figures 1C/F & 2, red)**. Additionally, a novel furin cleavage site has been reported in the SARS-CoV-2 virus, resulting in increased infectivity (Wrapp et al., 2020). Indeed, we find that the epitope SYQTQTNSPRRARSVASQSIIAYTMSLGAENSV containing the RRAR furin cleavage site of the spike protein ranks in the 90.7^th^ percentile of T cell epitopes and ranks first of 1546 in combined B and T cell epitope, (**Figures 1C/F & 2, orange)**, thereby targeting an additional evolutionary adaptation of SARS-CoV-2. Based on a recently published study identifying receptor binding hotspots deduced by comparing structures of ACE2 bound to the Spike protein from SARS-CoV-2 as compared to SARS-CoV (Shang et al., 2020), we searched for epitopes containing the five acquired residues that increase Spike binding to ACE2, identifying KPFERDISTEIYQ**<u>A</u>**GSTP**<u>C</u>**NGVEG**<u>FNC</u>**YFPLQS as the highest ranked epitope containing all of these residues (hotspots underlined; **Table 1**). Finally, it is known that mRNA transcripts proximal to the 3’ end of the *Coronaviridae* family genome show higher abundance consistent with the viral replication process, with S, E, M, and N genes shown to have significantly higher translational efficiency compared to the 5’ transcripts (Cheng, Lau, Woo, & Yuen, 2007; Hiscox, Cavanagh, & Britton, 1995; Irigoyen et al., 2016). We therefore posit that viral epitopes derived from 3’ terminus including the S, E, M, and N genes will have a higher representation on MHC and suggest their prioritization in a vaccine construct. Tables 1, Figure 2, Tables S2, and S3 show viral epitopes we suggest prioritizing for vaccine development.

**Figure 2.**
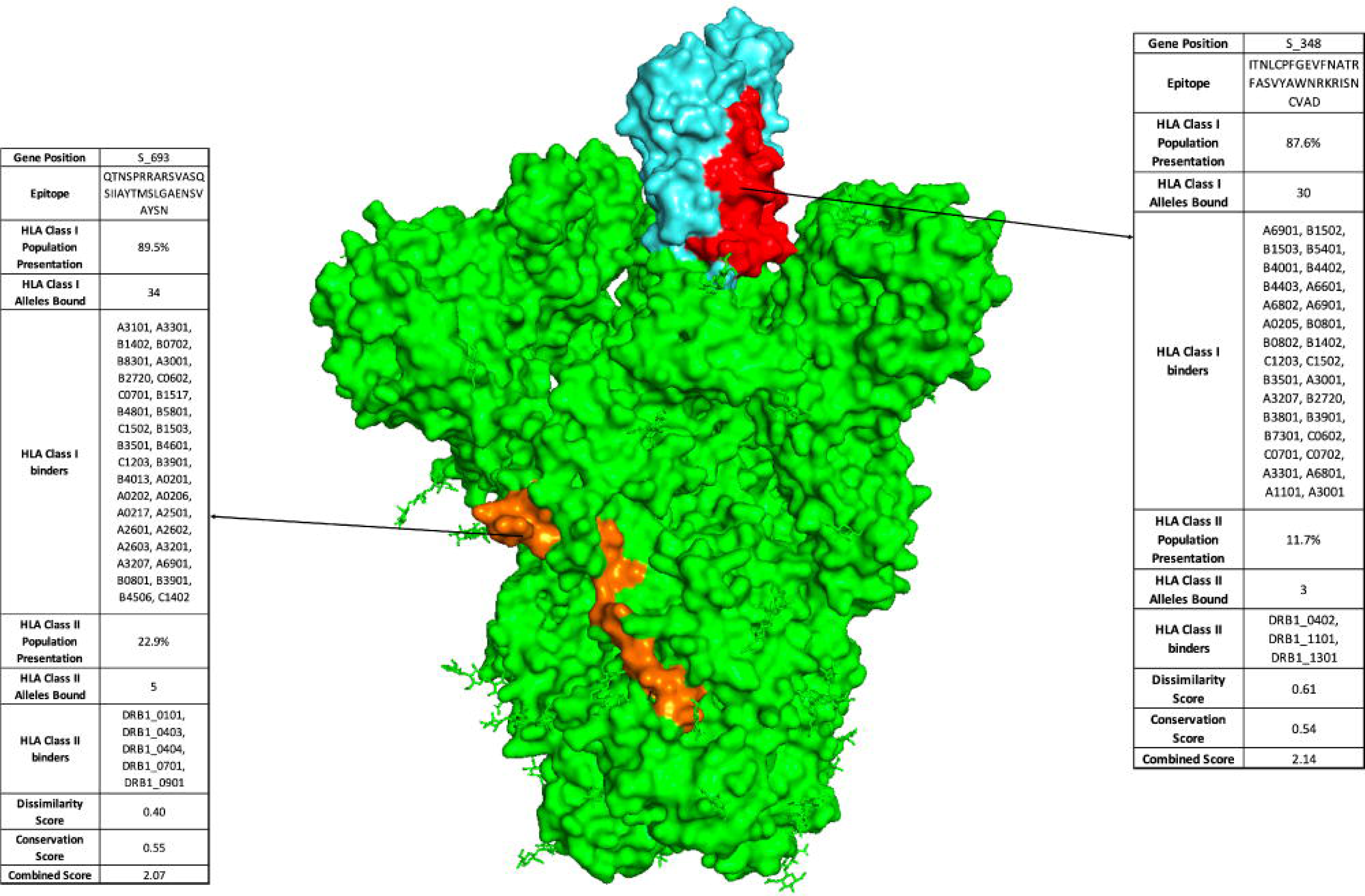
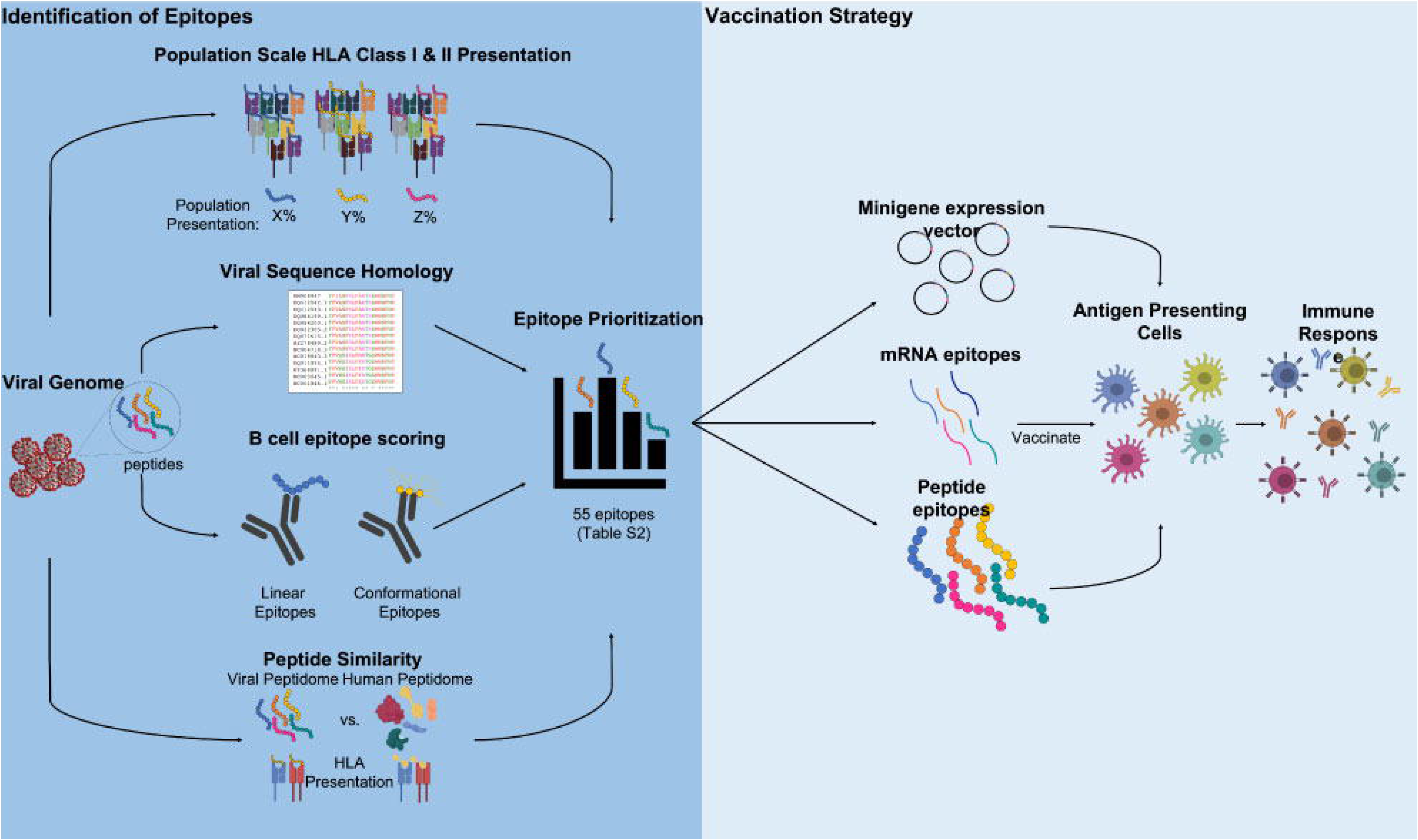
Proposed vaccine epitopes in SARS-CoV-2 Spike protein. Crystal structure of SARS-CoV-2 Spike protein trimer (PDB 6VYB) with two highlighted vaccine epitopes targeting novel acquired viral vulnerabilities. 1) SARS-CoV-2 receptor binding domain (cyan) has up to 10-fold higher affinity binding to the ACE2 receptor as compared to previous coronaviruses. Using our analysis, we identify a high-ranking vaccine epitope (red) within the receptor binding domain. 2) SARS-CoV-2 has acquired a novel furin cleavage site RRAR, along for increased infectivity due to improved membrane fusion (epitope containing the novel furin cleavage site highlighted in orange).

## Discussion

Here we present a comprehensive immunogenicity map of the SARS-CoV-2 virus (**Table S1**), highlighting 65 B and T cell epitopes (**Tables 1, S1 and S2**) from a diverse sampling of viral domains across all 9 SARS-CoV-2 genes. Based on our computational algorithm, we expect that the highest scoring peptides will result in safe and immunogenic T cell epitopes, and that B cell epitopes should be evaluated for safety and efficacy using methods previously reported (Wang et al., 2016). mRNA vaccines have been shown to be safe and effective in preclinical studies (Richner et al., 2017), with nucleoside RNAs shown to be effective without triggering RNA-induced immunogenicity (Pardi et al., 2017), while DNA vaccines have also been shown to be safe and protective (Dowd et al., 2016). Both DNA and mRNA vaccines are capable of being rapidly and efficiently manufactured at large scales. We suggest that a multivalent construct composed of the SARS-CoV-2 minigenes (presented in Tables 1, S2, and S3) can be used in a DNA on mRNA vaccine for expression in antigen-presenting cells. These epitopes can be used in tandem with a TLR agonist such as tetanus toxoid (Zanetti, Ferreira, de Vasconcelos, & Han, 2019) to drive activation of signals 1 and 2 in antigen presenting cells. Constructs can be designed to contain a combination of optimal B and T cell epitopes, or deployed as a construct consisting of the top scoring T cell epitopes to be used in combination with the vaccines currently being developed targeting the Spike protein in order to drive the adaptive memory response. DNA vaccine sequences can also be codon optimized to increase CpG islands such as to increase TLR9 activation (Krieg, 2008).

The methods described here provide a rapid workflow for evaluating and prioritizing safe and immunogenic regions of a viral genome for use in vaccination. With the third epidemic in the past two decades underway, and all originating from a coronavirus family virus, these viruses will continue to threaten the human population, and necessitate the need for prophylactic measures against future outbreaks. A subset of the epitopes selected here are derived from viral regions sharing a high degree of homology with other viruses in the family, and thus we expect these evolutionarily conserved regions to be essential in the infectivity and replicative lifecycle across the coronavirus family, suggesting that an immune response against the epitopes listed herein may provide more broadly protective immunity against other coronaviruses. Additionally, we describe epitopes containing the newly acquired features of SARS-CoV-2 that confer evolutionary advantages in viral spread and infectivity. In addition, the immunogenicity map provided in Table S1 can be used to customize epitopes based on the HLA frequencies of specific populations. Though here we suggest the use of 33mers based on optimal MHC presentation across the population, these methods can be applied to evaluate k-mers of various sizes depending on desired application.

Antigenic burden from epitopes that do not contribute to viral protection can cause autoimmune reactions, reactogenicity, detract from the efficacy of the virus, or result in ADE. To mitigate these effects *a priori*, we selected maximally immunogenic epitopes with the highest degree of dissimilarity to the self-proteome such as to minimize the potential of cross-reactivity that can lead to adverse reaction or minimize the efficacy of the virus. In addition to the predicted safety of these epitopes stemming from lack of potentially cross-reactive normal proteins, we expect that a greater repertoire of viral antigen-specific T cells will exist due to lack of negative thymic selection. We prioritize epitopes with maximal dissimilarity from the human proteome, however, many other SARS-CoV-2 peptides show identical or nearly identical peptides presented on MHC derived from normal proteins, suggesting their use in vaccination could result in an autoimmune response. The 65 epitopes presented here can be expressed in a ∼6.3kb construct and coupled with the safe and rapid production of synthetic DNA, mRNA, and peptide vaccines. As SARS-CoV-2 has precipitated the need to develop novel approaches to rapidly deploy vaccines in pandemic situations (Lurie, Saville, Hatchett, & Halton, 2020), we suggest that this comprehensive analysis can be incorporated into a process that can be rapidly deployed when future novel viral pathogens emerge.

## STAR Methods

### Population-scale HLA Class I & II Presentation

We identified potential SARS-CoV-2 epitopes by applying our recently published algorithm for scoring population-scale HLA presentation of tumor driver gene, to the SARS-CoV-2 genome (GenBank Acc#: MN908947.3) (Yarmarkovich et al., 2020). All possible 33mer amino acid sequences covering every 9mer peptide from the 10 SARS-CoV-2 genes were generated and we employed netMHC-4.0 to predict the binding affinities of each viral peptide across 84 HLA class I alleles. We considered peptides with binding affinities <500nM putative epitopes. MHC class II binding affinities were predicted as previously described across 36 HLA class II alleles population using netMHCII 2.3.

The frequencies of HLA class I alleles -A/B/C and HLA class II alleles -DRB1/3/4/5 were obtained from Be the Match bone marrow registry (Gragert et al., 2013).HLA class II alleles -DQA1/DQB1 and -DPA1/DPB1 were obtained from (Sidney et al., 2010) and (Solberg et al., 2008), respectively.

### Conservation Scoring

We obtained all 1,024 unique protein sequences categorized by each of the 10 SARS-CoV-2 genes available from the NCBI as of 25 March 2020. All sequences were aligned using Clustal Omega (Sievers et al., 2011) and each position summed for homology. In addition to human sequences, we scored each amino acid position for homology across 15 species of related coronavirus found in bats, pigs, camels, mice, and humans (SARS-CoV, SARS-CoV-2, and MERS). Each amino acid was scored up to 100% conservation. 33mer peptides were then scored in Equation 1:

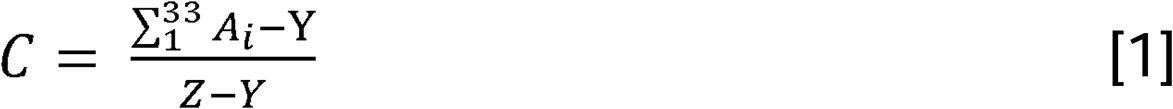

Where C is the 33mer conservation score, A is the conservation percentage of an amino acid position, Y is the minimum 33mer conservation percentage sum, and Z is the maximum 33mer conservation percentage sum. In the same way, we ranked the conservation across 274 SARS-CoV-2 amino acid sequences available at the time of this study. A final conservation score was generated by averaging the conservation scores from cross-species and interhuman variation and 33mer peptides with the highest score were considered the most conserved.

### Dissimilarity Scoring

3,524 viral epitopes were compared against the normal human proteome on each of their MHC binding partners, testing a total of 12, 383 peptide/MHC pairs against the entire human proteome (85,915,364 normal peptides across HLAs), assigning a similarity score for each peptide. Residues in the same position of the viral and human peptides with a perfect match, similar amino acid classification, or different polarity, were assigned scores of five, two, or negative two respectively. Similarity scores were calculated based on amino acid classification and hydrophobicity were determined using non-anchor residues of MHC (**Figure S1A**). The canonical TCR-interaction hotspots (residues four through six) were double weighted (Gagnon et al., 2005; Gras et al., 2009; Ishizuka et al., 2008). The similarity scores generated for each viral peptide were converted to Z-scores and peptides with a p <0.0001 were selected for comparison to viral epitopes (**Figure S1B**). The overall dissimilarity score for the viral peptide was then calculated using Equation 2:

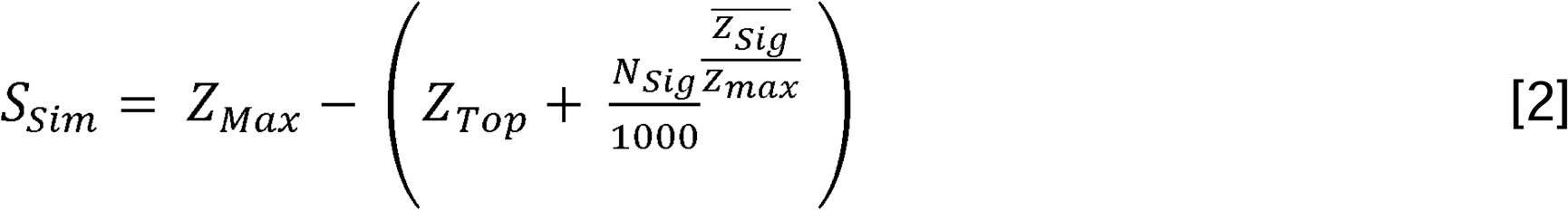

where *S*_*Sim*_ is the overall dissimilarity score for the viral peptide, *Z*_*Max*_ is the highest possible Z-score given a perfect sequence match to the viral peptide, *Z*_*Top*_ is the highest Z-score from the human proteome, *N*_*Sig*_ is the number of statistically significant peptides from the human proteome, and 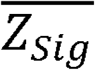 is the mean Z-score from the statistically significant peptides given a p < 0.001.

### B cell Epitope Scoring

We used BepiPred 2.0 and DiscoTope 2.0 (Jespersen et al., 2017; Kringelum et al., 2012) to score individual amino acid residues, assessing linear epitopes in Matrix, Envelope, and Spike proteins, and conformational epitopes for Spike protein, based on published structure (PDB 6VYB). To we summed and normalized linear and conformational, using separate normalizations for proteins in which only linear predictions were available.

## Author Contributions

Conceptualization, M.Y.; Methodology, M.Y., J.M.W, A.F.; Software, M.Y. J.M.W., and A.F.; Formal Analysis, M.Y., J.M.W., and A.F.; Investigation, M.Y., J.M.W., and A.F.; Funding Acquisition, M.Y. and J.M.M., Writing, M.Y., J.M.W, and J.M.M.; Supervision, M.Y. and J.M.M.

## Acknowledgements

This work was supported by a St. Baldrick’s-Stand Up To Cancer Pediatric Dream Team Translational Research Grant (SU2C-AACR-DT2727) and the Beau Biden Cancer Moonshot Pediatric Immunotherapy Discovery and Development Networ (NCI Grant U54 CA232568). Stand Up To Cancer is a program of the Entertainment Industry Foundation administered by the American Association for Cancer Research. This work was also supported by NIH R35 CA220500 and the Giulio D’Angio Endowed Chair and the Quod Erat Demonstrandum (QED) program at the Science Center in Philadelphia.

---

**Figure S1.**
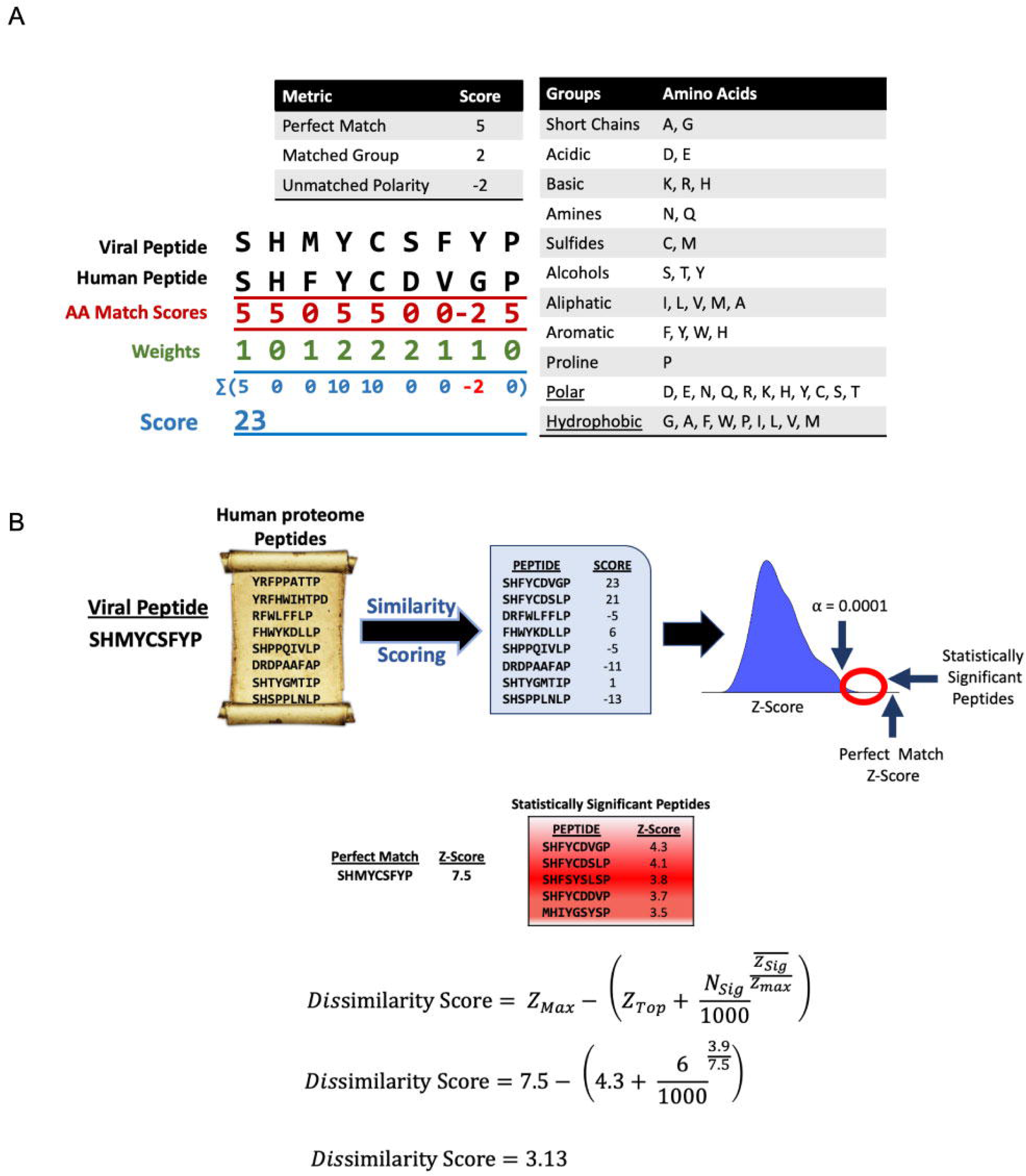
Dissimilarity Scoring. A) 3,524 viral epitopes (12,383 total peptide/MHC pairs) were compared against the normal human proteome. Non-anchor residues were used to calculate similarity scores based on amino acid classifications as described in Methods. Residues in the same position of the viral and human peptides with a perfect match, similar amino acid classification, or different polarity, were assigned scores of five, two, or negative two respectively. B) Each viral peptide/HLA pair was compared against the set of normal peptides presented on the same MHC. Dissimilarity score for each viral peptide calculated by comparing against the most similar group of peptides with p<0.0001, and reported as the difference in Z-scores between the viral peptide and closest-scoring peptides.

## References

Alspach, E., Lussier, D. M., Miceli, A. P., Kizhvatov, I., DuPage, M., Luoma, A. M., … Schreiber, R. D. (2019). MHC-II neoantigens shape tumour immunity and response to immunotherapy. Nature, 574(7780), 696–701. doi: 10.1038/s41586-019-1671-8

An, L.-L., Rodriguez, F., Harkins, S., Zhang, J., & Whitton, J. L. (2000). Quantitative and qualitative analyses of the immune responses induced by a multivalent minigene DNA vaccine. Vaccine, 18(20), 2132–2141. doi: https://doi.org/10.1016/S0264-410X(99)00546-0

Chen, J., Lee, K. H., Steinhauer, D. A., Stevens, D. J., Skehel, J. J., & Wiley, D. C. (1998). Structure of the Hemagglutinin Precursor Cleavage Site, a Determinant of Influenza Pathogenicity and the Origin of the Labile Conformation. Cell, 95(3), 409–417. doi: 10.1016/S0092-8674(00)81771-7

Cheng, V. C. C., Lau, S. K. P., Woo, P. C. Y., & Yuen, K. Y. (2007). Severe Acute Respiratory Syndrome Coronavirus as an Agent of Emerging and Reemerging Infection. Clinical Microbiology Reviews, 20(4), 660–694. doi: 10.1128/cmr.00023-07

Dejnirattisai, W., Supasa, P., Wongwiwat, W., Rouvinski, A., Barba-Spaeth, G., Duangchinda, T., … Screaton, G. R. (2016). Dengue virus sero-cross-reactivity drives antibody-dependent enhancement of infection with zika virus. Nature Immunology, 17(9), 1102–1108. doi: 10.1038/ni.3515

Dowd, K. A., Ko, S.-Y., Morabito, K. M., Yang, E. S., Pelc, R. S., DeMaso, C. R., … Graham, B. S. (2016). Rapid development of a DNA vaccine for Zika virus. Science, 354(6309), 237–240. doi: 10.1126/science.aai9137

Gagnon, S. J., Borbulevych, O. Y., Davis-Harrison, R. L., Baxter, T. K., Clemens, J. R., Armstrong, K. M., … Baker, B. M. (2005). Unraveling a Hotspot for TCR Recognition on HLA-A2: Evidence Against the Existence of Peptide-independent TCR Binding Determinants. Journal of Molecular Biology, 353(3), 556–573. doi: https://doi.org/10.1016/j.jmb.2005.08.024

Gragert, L., Madbouly, A., Freeman, J., & Maiers, M. (2013). Six-locus high resolution HLA haplotype frequencies derived from mixed-resolution DNA typing for the entire US donor registry. Hum Immunol, 74(10), 1313–1320. doi: 10.1016/j.humimm.2013.06.025

Gras, S., Saulquin, X., Reiser, J.-B., Debeaupuis, E., Echasserieau, K., Kissenpfennig, A., … Housset, D. (2009). Structural Bases for the Affinity-Driven Selection of a Public TCR against a Dominant Human Cytomegalovirus Epitope. The Journal of Immunology, 183(1), 430–437. doi: 10.4049/jimmunol.0900556

Hiscox, J. A., Cavanagh, D., & Britton, P. (1995). Quantification of individual subgenomic mRNA species during replication of the coronavirus transmissible gastroenteritis virus. Virus Research, 36(2), 119–130. doi: https://doi.org/10.1016/0168-1702(94)00108-O

Irigoyen, N., Firth, A. E., Jones, J. D., Chung, B. Y. W., Siddell, S. G., & Brierley, I. (2016). High-Resolution Analysis of Coronavirus Gene Expression by RNA Sequencing and Ribosome Profiling. PLoS pathogens, 12(2), e1005473–e1005473. doi: 10.1371/journal.ppat.1005473

Ishizuka, J., Stewart-Jones, G. B. E., van der Merwe, A., Bell, J. I., McMichael, A. J., & Jones, E. Y. (2008). The Structural Dynamics and Energetics of an Immunodominant T Cell Receptor Are Programmed by Its Vβ Domain. Immunity, 28(2), 171–182. doi: 10.1016/j.immuni.2007.12.018

Jespersen, M. C., Peters, B., Nielsen, M., & Marcatili, P. (2017). BepiPred-2.0: improving sequence-based B-cell epitope prediction using conformational epitopes. Nucleic Acids Research, 45(W1), W24–W29. doi: 10.1093/nar/gkx346

Krieg, A. M. (2008). Toll-like receptor 9 (TLR9) agonists in the treatment of cancer. Oncogene, 27(2), 161–167. doi: 10.1038/sj.onc.1210911

Kringelum, J. V., Lundegaard, C., Lund, O., & Nielsen, M. (2012). Reliable B Cell Epitope Predictions: Impacts of Method Development and Improved Benchmarking. PLOS Computational Biology, 8(12), e1002829. doi: 10.1371/journal.pcbi.1002829

Lu, Y.-C., Yao, X., Crystal, J. S., Li, Y. F., El-Gamil, M., Gross, C., … Robbins, P. F. (2014). Efficient identification of mutated cancer antigens recognized by T cells associated with durable tumor regressions. Clinical cancer research : an official journal of the American Association for Cancer Research, 20(13), 3401–3410. doi: 10.1158/1078-0432.CCR-14-0433

Lurie, N., Saville, M., Hatchett, R., & Halton, J. (2020). Developing Covid-19 Vaccines at Pandemic Speed. New England Journal of Medicine. doi: 10.1056/NEJMp2005630

McHeyzer-Williams, M., Okitsu, S., Wang, N., & McHeyzer-Williams, L. (2012). Molecular programming of B cell memory. Nature Reviews Immunology, 12(1), 24–34. doi: 10.1038/nri3128

Pardi, N., Hogan, M. J., Pelc, R. S., Muramatsu, H., Andersen, H., DeMaso, C. R., … Weissman, D. (2017). Zika virus protection by a single low-dose nucleoside-modified mRNA vaccination. Nature, 543(7644), 248–251. doi: 10.1038/nature21428

Richner, J. M., Himansu, S., Dowd, K. A., Butler, S. L., Salazar, V., Fox, J. M., … Diamond, M. S. (2017). Modified mRNA Vaccines Protect against Zika Virus Infection. Cell, 168(6), 1114–1125.e1110. doi: https://doi.org/10.1016/j.cell.2017.02.017

Shang, J., Ye, G., Shi, K., Wan, Y., Luo, C., Aihara, H., … Li, F. (2020). Structural basis of receptor recognition by SARS-CoV-2. Nature. doi: 10.1038/s41586-020-2179-y

Sidney, J., Steen, A., Moore, C., Ngo, S., Chung, J., Peters, B., & Sette, A. (2010). Divergent motifs but overlapping binding repertoires of six HLA-DQ molecules frequently expressed in the worldwide human population. J Immunol, 185(7), 4189–4198. doi: 10.4049/jimmunol.1001006

Sievers, F., Wilm, A., Dineen, D., Gibson, T. J., Karplus, K., Li, W., … Higgins, D. G. (2011). Fast, scalable generation of high-quality protein multiple sequence alignments using Clustal Omega. Mol Syst Biol, 7, 539. doi: 10.1038/msb.2011.75

Solberg, O. D., Mack, S. J., Lancaster, A. K., Single, R. M., Tsai, Y., Sanchez-Mazas, A., & Thomson, G. (2008). Balancing selection and heterogeneity across the classical human leukocyte antigen loci: a meta-analytic review of 497 population studies. Hum Immunol, 69(7), 443–464. doi: 10.1016/j.humimm.2008.05.001

Tetro, J. A. (2020). Is COVID-19 receiving ADE from other coronaviruses? Microbes and Infection, 22(2), 72–73. doi: https://doi.org/10.1016/j.micinf.2020.02.006

Thevarajan, I., Nguyen, T. H. O., Koutsakos, M., Druce, J., Caly, L., van de Sandt, C. E., … Kedzierska, K. (2020). Breadth of concomitant immune responses prior to patient recovery: a case report of non-severe COVID-19. Nature Medicine. doi: 10.1038/s41591-020-0819-2

Walls, A. C., Park, Y.-J., Tortorici, M. A., Wall, A., McGuire, A. T., & Veesler, D. Structure, Function, and Antigenicity of the SARS-CoV-2 Spike Glycoprotein. Cell. doi: 10.1016/j.cell.2020.02.058

Wan, Y., Shang, J., Sun, S., Tai, W., Chen, J., Geng, Q., … Li, F. (2020). Molecular Mechanism for Antibody-Dependent Enhancement of Coronavirus Entry. Journal of Virology, 94(5), e02015–02019. doi: 10.1128/jvi.02015-19

Wang, Q., Zhang, L., Kuwahara, K., Li, L., Liu, Z., Li, T., … Liu, G. (2016). Immunodominant SARS Coronavirus Epitopes in Humans Elicited both Enhancing and Neutralizing Effects on Infection in Non-human Primates. ACS infectious diseases, 2(5), 361–376. doi: 10.1021/acsinfecdis.6b00006

Wrapp, D., Wang, N., Corbett, K. S., Goldsmith, J. A., Hsieh, C.-L., Abiona, O., … McLellan, J. S. (2020). Cryo-EM structure of the 2019-nCoV spike in the prefusion conformation. Science, 367(6483), 1260–1263. doi: 10.1126/science.abb2507

Xu, H., Zhong, L., Deng, J., Peng, J., Dan, H., Zeng, X., … Chen, Q. (2020). High expression of ACE2 receptor of 2019-nCoV on the epithelial cells of oral mucosa. International Journal of Oral Science, 12(1), 8. doi: 10.1038/s41368-020-0074-x

Yarmarkovich, M., Farrel, A., Sison, A., di Marco, M., Raman, P., Parris, J. L., … Maris, J. M. (2020). Immunogenicity and Immune Silence in Human Cancer. Frontiers in Immunology, 11(69). doi: 10.3389/fimmu.2020.00069

Zanetti, B. F., Ferreira, C. P., de Vasconcelos, J. R. C., & Han, S. W. (2019). scFv6.C4 DNA vaccine with fragment C of Tetanus toxin increases protective immunity against CEA-expressing tumor. Gene Therapy, 26(10), 441–454. doi: 10.1038/s41434-019-0062-y

Zhang, H., Zhou, P., Wei, Y., Yue, H., Wang, Y., Hu, M., … Du, R. (2020). Histopathologic Changes and SARS–CoV-2 Immunostaining in the Lung of a Patient With COVID-19. Annals of Internal Medicine. doi: 10.7326/m20-0533

